# Itaconate and its derivatives repress C2C12 myogenesis

**DOI:** 10.1101/2021.02.01.429228

**Authors:** Tae Seok Oh, Damian C. Hutchins, Rabina Mainali, Kevin Goslen, Matthew A. Quinn

**Affiliations:** Department of Pathology, Section on Comparative Medicine, Winston-Salem, North Carolina 27517, USA; Department of Internal Medicine, Section on Molecular Medicine Wake Forest School of Medicine, Winston-Salem, North Carolina 27517, USA

**Keywords:** C2C12, myogenesis, itaconate, succinate, succinate dehydrogenase

## Abstract

A Krebs cycle intermediate metabolite, itaconate, has gained attention as a potential antimicrobial and autoimmune disease treatment due to its anti-inflammatory effects. While itaconate and its derivatives pose an attractive therapeutic option for the treatment of inflammatory diseases, the effects outside the immune system still remain limited, particularly in the muscle. Therefore, we endeavored to determine if itaconate signaling impacts muscle differentiation. Utilizing the well-established C2C12 model of *in vitro* myogenesis, we evaluated the effects of itaconate and its derivatives on transcriptional and protein markers of muscle differentiation as well as mitochondrial function. We found itaconate and the derivatives dimethyl itaconate and 4-octyl itaconate disrupt differentiation media-induced myogenesis. A primary biological effect of itaconate is a succinate dehydrogenase (SDH) inhibitor. We find the SDH inhibitors dimethyl malonate and harzianopyridone phenocopie the anti-myogenic effects of itaconate. Furthermore, we find treatment with exogenous succinate results in blunted myogenesis. Together our data indicate itaconate and its derivatives interfere with *in vitro* myogenesis, potentially through inhibition of SDH and subsequent succinate accumulation. More importantly, our findings suggest the therapeutic potential of itaconate and its derivatives could be limited due to deleterious effects on myogenesis.

## Introduction

It is well appreciated that dramatic metabolic reprogramming occurs in response to inflammatory challenges. One of the most robust metabolic adaptations made in myeloid cells in response to inflammatory cues is the diversion of cis-aconitate away from α-ketoglutarate towards the synthesis of itaconate (1,2). One primary function itaconate serves is to reduce inflammation via the inhibition of succinate dehydrogenase (SDH) thereby reducing accumulation of reactive oxygen species (ROS) (1). Due to its potent anti-inflammatory properties, itaconate and its derivatives such as dimethyl itaconate (DMI) and 4-octyl itaconate (4-OI) have gained attention as potential treatments for autoimmune conditions such as psoriasis (3), multiple sclerosis (4), rheumatoid arthritis (5), and systemic lupus erythematosus (6).

While the signaling attributes of itaconate with regards to immune function are well delineated, not much is known on the actions of itaconate in non-immune organs. What is known is that itaconate possesses anti-inflammatory, anti-fibrotic and pro-survival attributes in organs such as the liver (7) and kidney (8). However, the effects of itaconate regulation of muscle function is yet unknown.

Given that chronic inflammation is hypothesized to incur sustained itaconate production, coupled to its potential therapeutic use for the treatment of a variety of diseases, it is of the upmost importance to understand potential off target tissues and effects in the face of elevated itaconate levels. To address this critical gap in knowledge we utilized the well-established C2C12 murine myoblast model system for *in vitro* myogenesis and studied the effects of itaconate and its derivatives on altering myoblast differentiation.

## Results

### Itaconate derivatives inhibits myogenesis in C2C12 cells

Past research has shown that Krebs cycle metabolites serve as signal transducers in the immune response within macrophages (9). We endeavored to evaluate how the presence of itaconate affects myogenic differentiation within C2C12 cells. Given its recent focus as a potential treatment for autoimmune conditions, the function of dimethyl itaconate (DMI) was first investigated. Gene expression markers for myogenesis (i.e. *Myh1*, *Myh2*, *Myh7*, *Tnnt1*, and *Tnnt3*) were measured in the presence of DMI versus vehicle over a 4-day differentiation protocol. As expected, vehicle group cells had a time-dependent increase in the transcription of myogenic genes (Fig. 1a). However, treatment with DMI significantly inhibited the induction of these myogenic markers during our differentiation protocol (Fig. 1a). Another analog of itaconate was also utilized since DMI cannot be endogenously converted back to intracellular itaconate (10). 4-octyl itaconate (4-OI) was subsequently used because it has been suggested as a possible treatment for autoimmune conditions, but unlike DMI, it can be endogenously converted to itaconate (10,11). We analyzed C2C12 cells treated with 4-OI over 4 days in the presence of differentiation media (DM). 4-OI significantly inhibited the induction of myogenic gene transcription as well (Fig. 1b).

**Fig. 1.**
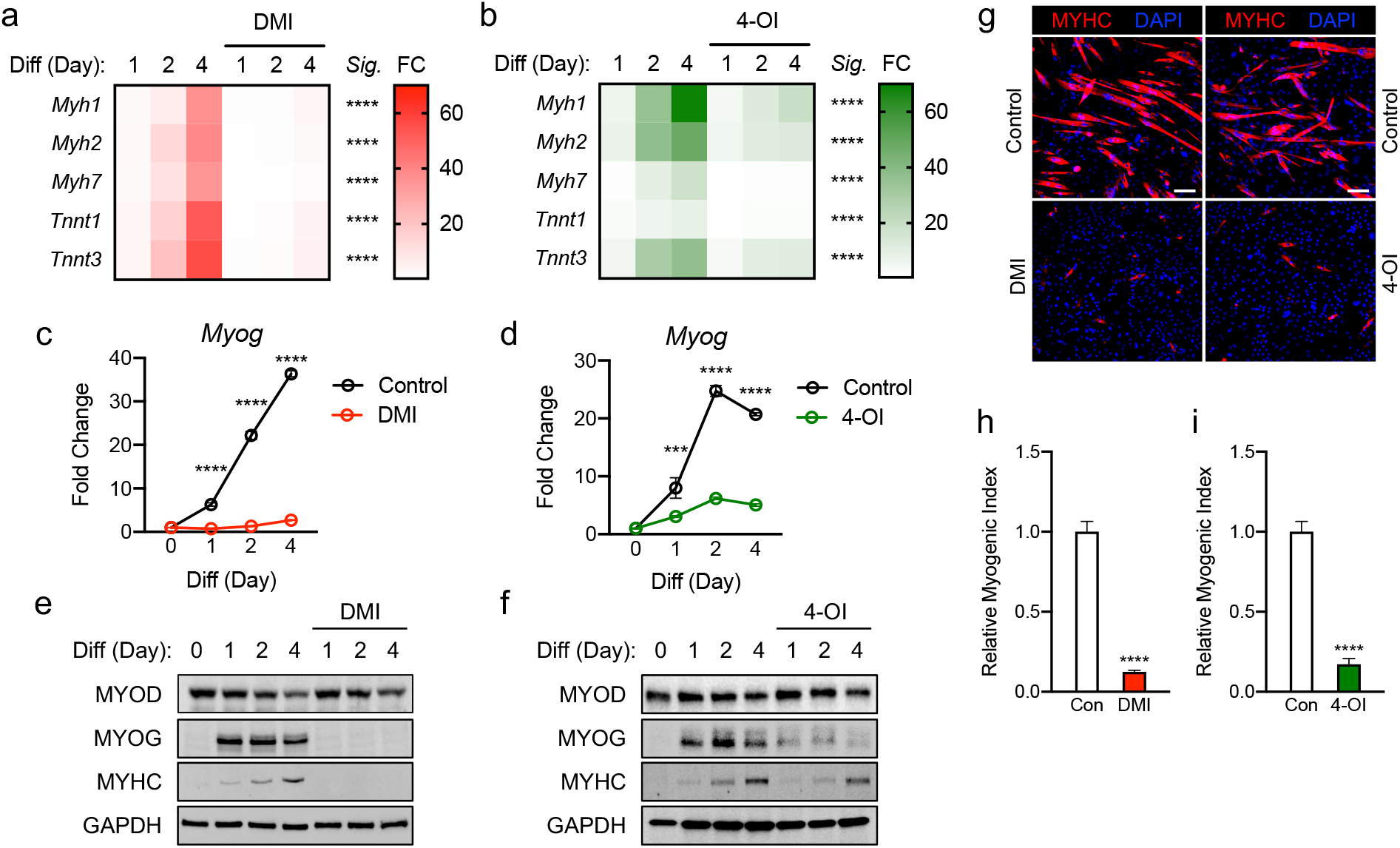
Itaconate derivatives represse C2C12 myogenesis by modulating myogenic transcription mechanisms. Heatmap depiction of average fold change (*Sig.* denotes statistical differences at day 4, n = 3) in myogenic gene transcription levels detected via RT-qPCR when exposed to differential media with or without (a) dimethyl itaconate (DMI: 125 μM) and (b) 4-octyl itaconate (4-OI: 125 μM) relative to the untreated group over a 4-day time course. Fold change of *Myog* mRNA expression at various time points comparing differentiation media (DM) control groups to (c) DMI and (d) 4-OI groups relative to expression without either treatment (n = 3). Western blotting data showing differences in myogenic transcription factor expression between DM treated groups and groups treated with DM plus (e) DMI and 4-OI over a 4 day time course as well as groups which were not exposed to either treatment for any period. (g) MYHC staining at day 4 of differentiation. Bar: 100 μm. Relative myogenic index (percentage of total nuclei associated with myotubes) of (h) DMI and (i) 4-OI. *** denotes *p* < 0.001, \*\*\*\**p* < 0.0001 based on two-way ANOVA/t-test.

To further determine how DMI and 4-OI impairs myogenesis, we next assessed the expression of the essential myogenic transcription factors *Myog* and *Myod.* The expression of *Myog* mRNA was significantly impaired by DMI throughout day 1 to day 4 of differentiation compared to the control (Fig. 1c). Transcription of *Myog* was also blunted in response to 4-OI exposure (Fig. 1d). Protein levels of MYOD were not different in the DMI groups compared to the control in response to DM. However, MYOG expression was almost completely abolished by DMI (Fig. 1e). Consequently, DMI-treated C2C12 myoblasts failed to differentiate into myotubes. DMI-treated groups showed no trace of MYHC in response to DM cues, while vehicle-treated cells displayed a time-dependent increase in MYHC protein (Fig. 1e). 4-OI significantly blocked MYOG and MYHC expression levels although it did not change MYOD levels (Fig. 1f). MYHC was visualized by immunofluorescence at day 4 of differentiation (Fig. 1g) and myogenic index was significantly impaired by DMI (Fig. 1h) and 4-OI (Fig. 1i).

### Physiological itaconate inhibits myogenesis

To investigate a role of physiological itaconate, C2C12 myoblast cells were then treated with DM only or DM plus itaconate and then examined over 4 days. It was reported that exogenous itaconate readily enters cells (9). Itaconate was used to portray physiological mechanisms more accurately. Transcription levels of *Myh2* and *Myh7* were not significantly impacted by itaconate exposure. However, the blunted induction of *Myh1*, *Tnnt1*, and *Tnnt3* was significant on the 4th day of exposure (Fig. 2a). Significant inhibition of *Myog* transcription on days 2 and 4 was also observed in itaconate groups (Fig. 2b). MYOG and MYHC expression were also inhibited in itaconate groups (Fig 2c). Itaconate treatment reduced MYHC staining (Fig. 2d) and myogenic index (Fig. 2e).

**Fig. 2.**
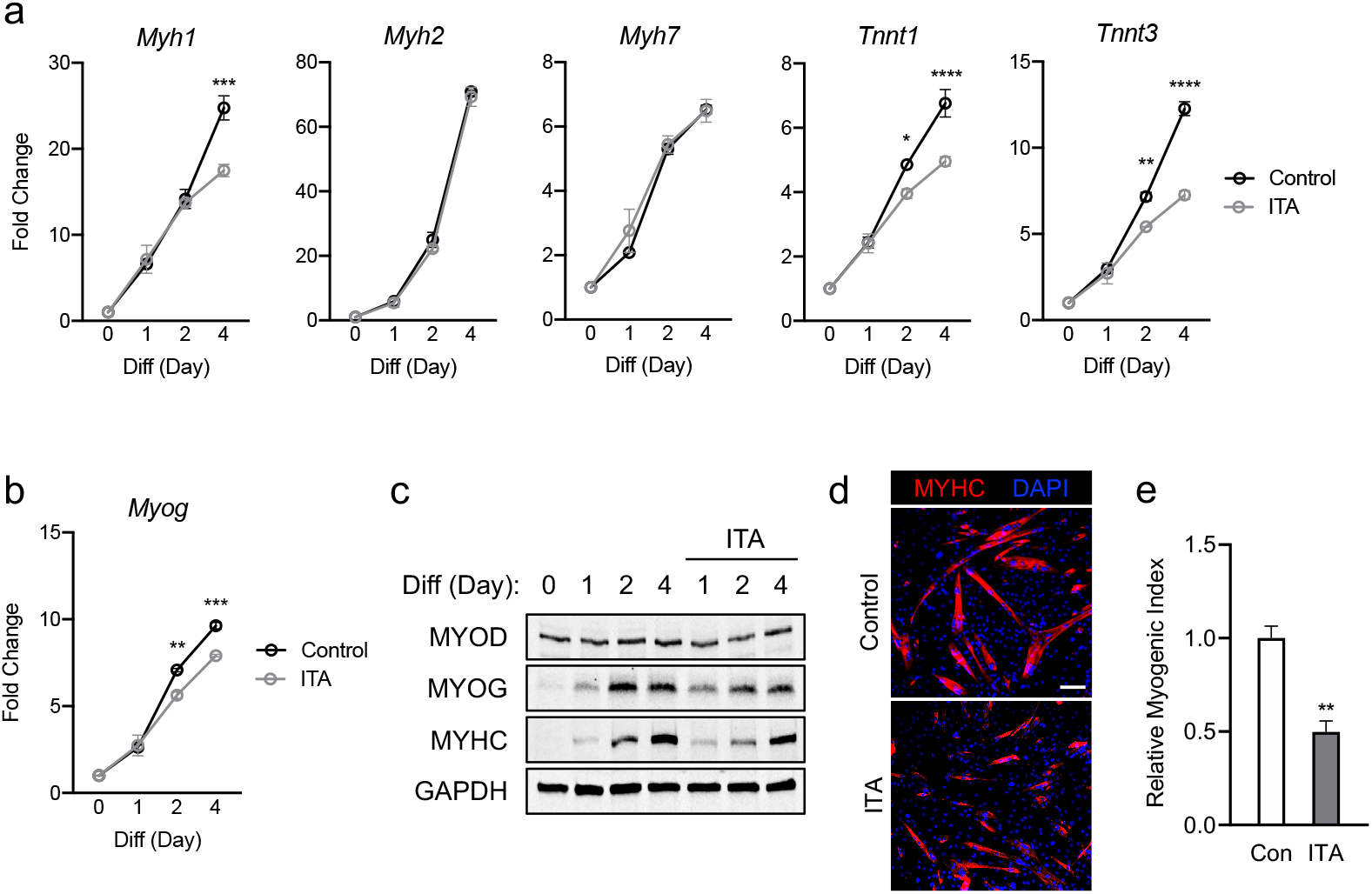
Itaconate represses C2C12 myogenesis by modulating myogenic transcription mechanisms. (a) 4-day time course showcasing fold change differences of myogenic gene transcription levels detected via RT-qPCR when exposed to differential media with or without itaconate (7.5 mM) relative to the untreated group (n = 3). (b) Fold change of *Myog* mRNA expression at various time points comparing DM control groups to DM and itaconate groups relative to expression without either treatment (n = 3). (c) Western blotting data showing differences in myogenic transcription factor expression between DM treated groups and groups treated with both DM and itaconate over a 4 day time course as well as groups which were not exposed to either treatment for any period. (d) MYHC staining at day 4 of differentiation. Bar: 100 μm. (e) Relative myogenic index (percentage of total nuclei associated with myotubes). * denotes *p* < 0.05, \*\**p* < 0.01, \*\*\**p* < 0.001, \*\*\*\**p* < 0.0001 based on two-way ANOVA/t-test.

Overall DMI shows the strongest myogenic inhibitory response and itaconate has the weakest amount of myogenic suppression among itaconate and derivatives tested. Taken together, these results suggest that itaconate and its derivatives are sufficient to inhibit myogenesis in C2C12 cells, modulating myogenic regulating factors at the transcription and protein levels.

### Malonate and pharmacological succinate dehydrogenase inhibitor obstruct myogenesis

Given that both itaconate and malonate are Krebs cycle metabolites that are shown to inhibit SDH (1,12), our next step was to determine whether malonate exposure could also inhibit C2C12 myogenesis. Dimethyl malonate (DMM) was administrated instead of itaconate under similar parameters to those discussed in Figures 1 through 2.

Malonate inhibited the induction of myogenic markers during myogenesis (Fig. 3a). This inhibition was more severe than itaconate but not as extreme as the inhibition caused by itaconate derivatives. In parallel, a potent and specific inhibitor of SDH, which is Harzianopyridone (Harz) possessing antibiotic and antifungal effects (13), was tested to determine whether this specific shared inhibition was the primary inducer of myogenesis repression. We examined C2C12 cells exposed to Harz using the same methods to elucidate whether SDH inhibition independent of the Krebs cycle might also elicit a dramatic obstruction of myogenesis. Harz treatment resulted in significant suppression of myogenic markers (Fig. 3b). Malonate lowered *Myog* levels compared to the control at all the time points (Fig. 3c). Consistent suppression of *Myog* expression was also observed following Harz treatment throughout the time course (Fig. 3d). MYOD levels were similar in malonate-treated cells during differentiation but malonate led to a reduction in MYOG levels, followed by a significant decrease in MYHC levels (Fig. 3e). Harz showed significantly reduced MYOG and MYHC levels as well (Fig. 3f). Malonate showed similar extent of reduction in MYHC staining (Fig. 3g) and myogenic index (Fig. 3h) as itaconate. Harz displayed the most robust inhibition of myogenesis shown by MYHC staining (Fig. 3g) and myogenic index (Fig. 3i). The degree of suppression caused by Harz was most comparable to the amount of inhibition by DMI. In tandem, these findings support that inhibition of SDH affects myogenesis in C2C12 cells.

**Fig. 3.**
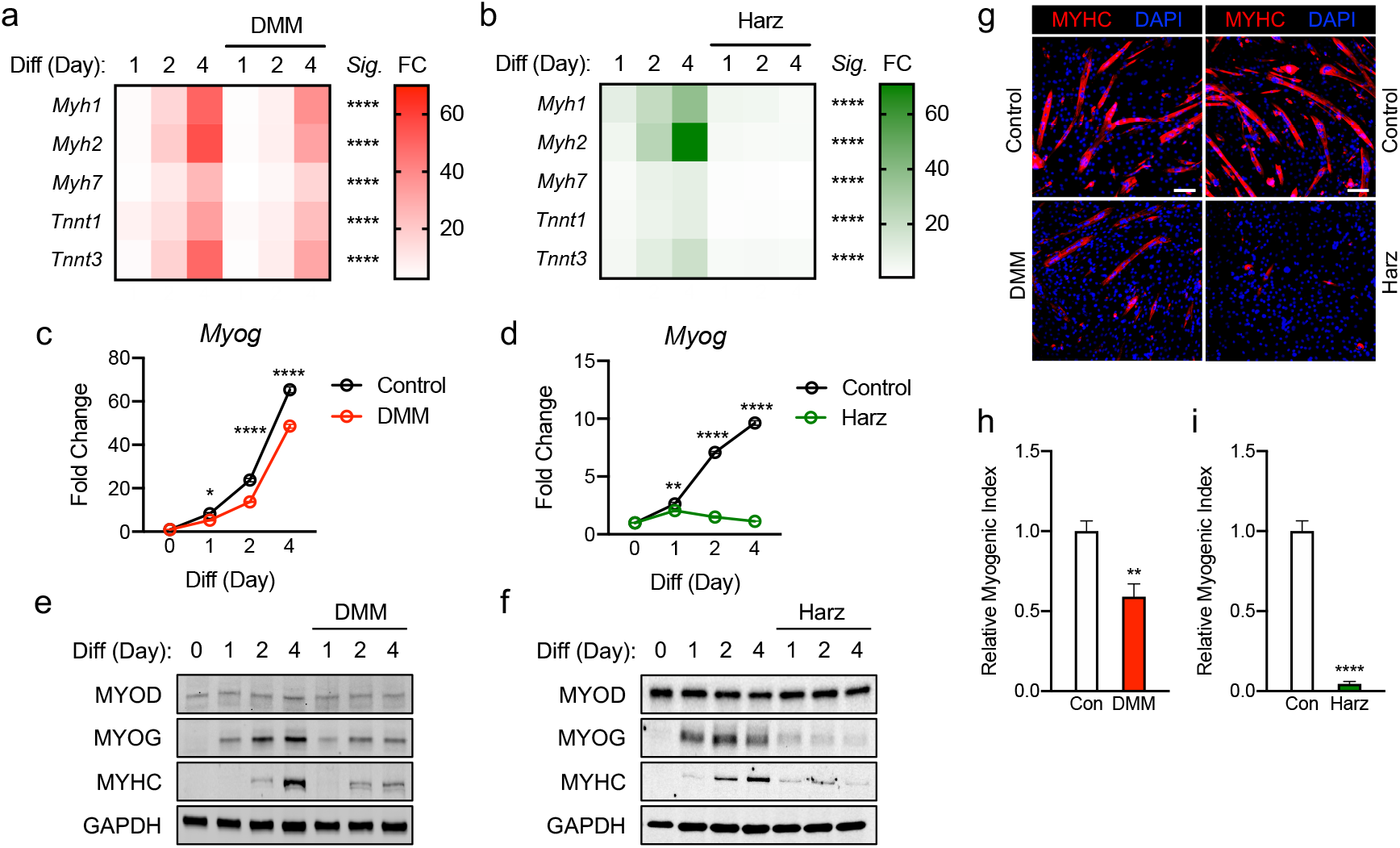
Malonate and pharmacological inhibitor of succinate dehydrogenase repress C2C12 myogenesis via similar mechanism of action to that of itaconate and its derivatives. Heatmap depiction of average fold change (*Sig.* denotes statistical differences at day 4, n = 3) in myogenic gene transcription levels detected via RT-qPCR when exposed to differential media with or without dimethyl malonate (DMM: 5 mM) and (b) harzianopyridone (Harz: 4 μM) relative to the untreated group over a 4-day time course. Fold change of *Myog* mRNA expression at various time points comparing differentiation media (DM) control groups to (c) DMI and (d) 4-OI groups relative to expression without either treatment (n = 3). Western blotting data showing differences in myogenic transcription factor expression between DM treated groups and groups treated with DM plus (e) DMM and (f) Harz over a 4 day time course as well as groups which were not exposed to either treatment for any period. (g) MYHC staining at day 4 of differentiation. Bar: 100 μm. Relative myogenic index (percentage of total nuclei associated with myotubes) of (h) DMM and (i) Harz. * denotes *p* < 0.05, \*\**p* < 0.01, \*\*\*\**p* < 0.0001 based on two-way ANOVA/t-test.

**Fig. 4.**
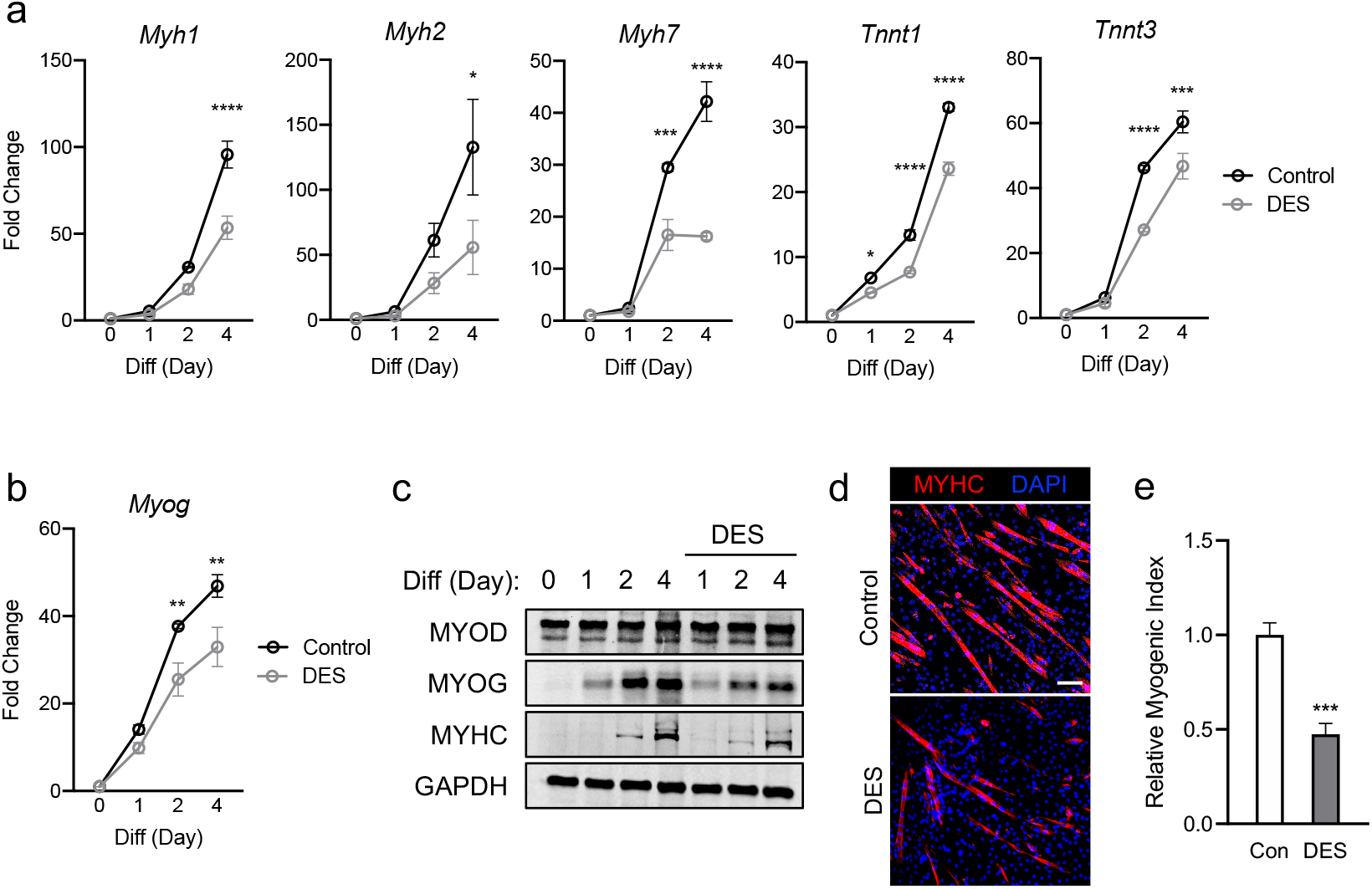
Elevated exposure to succinate elicits inhibition of C2C12 myogenesis via mechanism similar to those observed when exposing cells to itaconate and malonate. (a) 4-day time course showcasing fold change differences of myogenic gene transcription levels detected via RT-qPCR when exposed to differential media with or without diethyl succinate (DES: 5 mM) relative to the untreated group (n = 3). (b) Fold change of *Myog* expression at various time points comparing DM control groups to DM and DES groups relative to expression without either treatment (n = 3). (c) Western blotting data showing differences in myogenic transcription factor expression between DM treated groups and groups treated with both DM and DES over a 4 day time course as well as groups which were not exposed to either treatment for any period. (d) MYHC staining at day 4 of differentiation. Bar: 100 μm. (e) Relative myogenic index (percentage of total nuclei associated with myotubes). * denotes *p* < 0.05, \*\**p* < 0.01, \*\*\**p* < 0.001, \*\*\*\**p* < 0.0001 based on two-way ANOVA/t-test.

### Myogenesis is inhibited by exogenous succinate

Inhibition of SDH activity can lead to an increase in intracellular succinate levels (14). Given that SDH inhibition alone is able to significantly inhibit myogenesis, we next asked if an abundance of succinate could be the connecting factor leading to myogenic inhibition. Thus, we tested if exogenous succinate can cause myogenic failure in C2C12 cells treated with DM. All the data with diethyl succinate (DES) represented a very similar trend to previous findings. Myogenic gene expression (Fig. 6a), *Myog* (Fig. 6b), and myogenic proteins (Fig. 6c) were all inhibited by the exogenous succinate treatment. Reduced MYHC expression (Fig. 6d) and myogenic index (Fig. 6e) by succinate indicate its inhibitory potency of myogenesis. The degree of suppression was most similar to those results observed in groups treated with itaconate or malonate. These results support a possibility that inhibition of myogenesis by itaconate or malonate is at least partly due to accumulated succinate caused by SDH inhibition.

## Discussion

Itaconate has become the epitome of metabolites that also have immunomodulation properties. It is a decarboxylated cis-aconitate which is primarily known for its SDH inhibiting activity in the TCA cycle. While initially used as a polymer for industrial purposes, it has been researched to have both antimicrobial and anti-inflammatory properties. Some bacteria (e.g. *Yersinia pestis*) have itaconate degrading enzymes and itaconate itself has been observed to restrict the growth of bacteria by inhibiting isocitrate lyase activity (15). It also regulates host mechanisms. Its anti-inflammatory role has gained the most attention recently. Itaconate is observed to significantly increase in response to macrophage activation (12). It acts to reduce inflammation by inhibiting the release of proinflammatory cytokines through a plethora of mechanisms, inhibiting ROS production through suppression of SDH activity (1), activating the regulator of antioxidant expression NRF2 (16), and activating the anti-inflammatory signaling transcription factor ATF3 (3). These effects in tandem have attracted scientists to get approval for itaconate derivatives as potential treatments to limit microbial pathogenicity and to attenuate autoimmune disease symptoms.

DMI and 4-OI have been researched to have promising effects on psoriasis, rheumatoid arthritis, multiple sclerosis, and systemic lupus erythematosus due to their ability to suppress IL-17 signaling and reduce proinflammatory cytokine production (3,6). While seemingly propitious as a potential treatment, the fact is that knowledge is very limited on the effects of itaconate and its derivates outside of those relating to the immunomodulatory axis. A consequence of itaconate derivative use may be one or more harmful side effects which could lessen enthusiasm for their utilization as treatments. Blocking itaconate signaling, which has yet to be proposed, might also be used to attenuate symptoms of other conditions. The focus of our research is to showcase the additional effects of itaconate on muscle regulation. Chronic inflammation has been linked to muscle wasting. Prolonged itaconate exposure resulting from chronic inflammation may cause or exacerbate muscle wasting.

Our hypothesis is that itaconate obstructs myogenesis via inhibition of SDH. This hypothesis arose from research done on the effects of succinate abundance on muscle wasting. Succinate is a substrate for SDH, so its intracellular concentration is directly linked to SDH activity. Succinate abundance has been proposed as a causative agent of muscle wasting. Treatments to elevate intracellular succinate concentration has been observed to decrease total muscle protein expression by ~25%, inhibit in vitro C2C12 myoblast cell myogenesis, decrease myofiber diameter in murine models, inhibit myogenesis in vivo following barium chloride injury, reduce respiration capacity by ~35% in myoblasts, and also disrupt a number of other metabolic processes within muscle cells (17). SDH has also been shown to have significantly reduced activity in sarcopenic muscle (18). Given that anti-inflammatory activity of itaconate is incurred via SDH inhibition, we endeavored to determine if itaconate treatment led to similar effects seen in succinate abundant cells.

We first tested the proposed treatment forms of itaconate, DMI, and 4-OI. We then examined itaconate alone followed by malonate, a pharmacological inhibitor of SDH, and then lastly diethyl succinate. Treatment incurred transcriptional modulation of *Myh1, Myh2, Myh7, Tnnt1, and Tnnt3* along with alteration of myogenic transcription factor (mainly MYOG) and myogenesis marker MYHC protein expression were examined. Our results showcased substantial inhibition of myogenesis in C2C12 cells by DMI. 4-OI treatment also exhibited significant inhibition, but not as aggressive as DMI. Itaconate treatment also resulted in significant myogenesis inhibition, but not to the degree seen in either DMI or 4-OI treated groups. It is possible that inability of DMI conversion to itaconate leads to a strong inhibition of myogenesis exhibiting sustained activity because 4-OI can be converted to itaconate, but DMI cannot (10). It is established that itaconate can be converted to itaconyl-CoA by succinate-CoA ligase (19). A possible mechanism to explain this tiered order of inhibition severity may involve itaconate conversion to itaconyl-CoA. Regardless of potency, all itaconate treatments inhibited *in vitro* myogenesis.

We show that SDH suppression and subsequent succinate accumulation significantly inhibit myogenesis via myogenic transcription suppression. Given that MYOG expression is dependent upon MYOD activity, it is hypothesized that MYOD activity was also suppressed in the treatment groups, but via post translational modification that leads to changes in MYOD activity such as phosphorylation and acetylation (20,21) rather than transcriptional inhibition. Inactive MYOD in response to SDH inhibition and succinate causes blunted induction of *Myog* thereby suppressing the transcription of myogenic genes. Together, our results support that itaconate may contribute to impairment of myogenesis via its effects on SDH which modulates myogenic transcription mechanisms.

Sarcopenia patients have been observed to have inhibited mitochondrial functioning via transcriptional downregulation of key genes involved in oxidative phosphorylation and mitochondria proteostasis (18). 4-OI has been shown to increase aerobic respiration by reducing glycolytic function through inhibition of GAPDH (11). This data may implicate that in addition to subsequent muscle wasting induced by increased succinate concentration, prolonged itaconate activity may exacerbate issues relating to mitochondrial function. What demands further study is the determination of whether itaconate and its derivatives induce mitochondrial activity suppression which may cause subsequent inhibition of the oxidative phosphorylation pathway and thus reduce energy demand accommodation capability.

In this study, we find that itaconate and derivatives contribute to suppression of myogenesis. What remains to be researched is how itaconate affects muscle regulation outside of succinate accumulation. Our results showcase functions of itaconate outside of immunomodulation and implicate that derivative use to treat autoimmune diseases or microbial infections should be cautioned due to a potential for deleterious effects on myogenesis. Further elucidation of these functions may lead to the development of treatments to attenuate muscle wasting via possible itaconate signaling inhibition.

## Materials and methods

### Antibodies and chemical reagents

Primary antibodies used in this study are as follows: GAPDH (Santa Cruz Biotechnology, sc-32233), MYHC (Sigma, 05-716), MYOD (Santa Cruz Biotechnology, sc-377460), and MYOG (Santa Cruz Biotechnology, sc-12732). Reagents added to the media are as follows: Diethyl succinate (Sigma, 112402), Dimethyl itaconate (Sigma, 592498), Dimethyl malonate (Sigma 136441), Harzianopyridone (Santa Cruz Biotechnology, sc-280769), Itaconic acid (Sigma, I29204), and 4-octyl itaconate (Tocris Bioscience, 6662).

### Cell culture and differentiation

The mouse myogenic C2C12 myoblasts were maintained on plastic cell culture plates in Dulbecco’s modified Eagle’s medium (DMEM) supplemented with 10% fetal bovine serum and 1% penicillin-streptomycin in a humidified incubator kept at 37 °C and 5% CO_2_. Cells were used up until passage 9. For differentiation, cells at 90% confluency were serum restricted with differentiation medium (DMEM, 2% horse serum, 1% penicillin-streptomycin) for up to 4 days. Growth or differentiation medium was replenished every day.

### RNA isolation and RT-qPCR

RNA isolation was performed using the commercially available Aurum RNA miniprep kit (Bio-rad, 732-6820). Gene expression analysis was conducted with 50 ng RNA using iTaq Universal One-Step RT-qPCR Kit (Bio-rad, 1725140). The reaction was carried out according to the manufacturer’s instructions using CFX Connect Real-Time PCR Detection System (Bio-rad, 1855200). Probes used for TaqMan^®^ Gene Expression Assays (ThermoFisher) were as follows: *Myog* (Mm00446194_m1), *Myh1* (Mm01332489_m1), *Myh2* (Mm01332564_m1), *Myh7* (Mm00600555_m1), *Ppib* (Mm00478295_m1), *Tnnt1* (Mm00449089_m1), and *Tnnt3* (Mm01137842_g1).

### Western blot

Cells were lysed in RIPA buffer (ThermoFisher, 89901) in the presence of Halt™ Protease and Phosphatase Inhibitor Cocktail (ThermoFisher, 78440), and protein concentration was measured using Pierce™ BCA Protein Assay Kit (ThermoFisher, 23227). Lysates were boiled for 5 min with 4x Laemmli Sample Buffer (Bio-Rad, 1610747) and 2-mercaptoethanol (Bio-Rad, 1610710). Lysates were resolved on SDS polyacrylamide gels and blotted onto PVDF membranes using Trans-Blot Turbo RTA Midi 0.45 μm LF PVDF Transfer Kit (Bio-Rad, 1704275). Transfer was run in Trans-Blot Turbo Transfer System (Bio-Rad, 1704150) using Mixed MW (1.3A-25V-7M) protocol. The membranes were blocked with Intercept™ (TBS) Blocking Buffer (LI-COR, 927-60001) for 1 h at room temperature and incubated with appropriate primary antibodies diluted 1:1000 in blocking buffer at 4 °C overnight. After 3 washes with 0.05% tween20 (Bio-Rad) in TBS buffer (Bio-Rad), the membrane was incubated with IRDye^®^ 800CW Goat anti-Mouse IgG (LI-COR, 926-32210) and IRDye^®^ 680RD Goat anti-Rabbit IgG (LI-COR, 926-68071) secondary antibodies diluted 1:10000 in blocking buffer for 1 h at room temperature. Band images were visualized using ChemiDoc™ MP Imaging System (Bio-Rad, 12003154).

### Immunofluorescence and calculation of myogenic index

Cells were seeded in a 24-well cell culture plate and followed by above-mentioned differentiation protocol. At day 4 of differentiation, cells were rinsed with PBS and fixed with 4% paraformaldehyde for 10 min at room temperature. Then, cells were permeablized with 0.5% Triton™ X-100 (Fisher, BP151-100) in PBS for 10 min at room temperature. Next, cells were submerged with Intercept^®^ (PBS) Blocking Buffer (LI-COR, 927-70001) for 1 h at room temperature. MYHC primary antibody was diluted to 1:100 in blocking buffer and incubated overnight at 4 °C with the fixed cells. Next day, after 3 washes with 0.05% tween20 in PBS, Alexa Fluor 594 secondary antibody (Fisher, A11005) was diluted to 1:500 in blocking buffer and incubated with the cells for 1 h at room temperature. Followed by further washes, cells were mounted with VECTASHIELD^®^ Antifade Mounting Medium with DAPI (Vectalabs, H-1200) and covered with circular cover glass. Images were taken using ZOE™ Fluorescent Cell Imager (Bio-rad, 1450031). Myogenic index was calculated as the percentage of nuclei in fused myotubes (MYHC positive) out of the total nuclei in given images. Distribution of nuclei in myoblasts and myotubes was measured by counting over 100 nuclei at 4 distinct locations. The number of nuclei was counted using imageJ.

### Statistical analysis

Unpaired t-test or two-way ANOVA with multiple comparisons was performed to determine significant relationships between groups in the experiments. Data are represented as mean ± SEM.

## Notes

### Competing Interest Statement

The authors have declared no competing interest.

